# Microphthalmia and disrupted retinal development due to a *LacZ* knock-in/knock-out allele at the *Vsx2* locus

**DOI:** 10.1101/2024.06.08.597937

**Authors:** Francesca R. Napoli, Xiaodong Li, Alan A. Hurtado, Edward M. Levine

## Abstract

Visual System Homeobox 2 (*Vsx2*) is a transcription factor expressed in the developing retina that regulates tissue identity, growth, and fate determination. Several mutations in the *Vsx2* gene exist in mice, including a spontaneous nonsense mutation and two targeted missense mutations originally identified in humans. Here, we expand the genetic repertoire to include a *LacZ* reporter allele (*Vsx2^LacZ^*) designed to express beta-Galactosidase (b-GAL) and simultaneously disrupt *Vsx2* function (knock-in/knock-out). The retinal expression pattern of b-GAL is concordant with VSX2, and the mutant allele is recessive. *Vsx2^LacZ^* homozygous mice have congenital bilateral microphthalmia accompanied by defects in retinal development including ectopic expression of non-retinal genes, reduced proliferation, delayed neurogenesis, aberrant tissue morphology, and an absence of bipolar interneurons - all hallmarks of *Vsx2* loss-of-function. Unexpectedly, the mutant VSX2 protein is stably expressed, and there are subtle differences in eye size and early retinal neurogenesis when compared to the null mutant, *ocular retardation J*. The perdurance of the mutant VSX2 protein combined with subtle deviations from the null phenotype leaves open the possibility that *Vsx2^LacZ^* allele is not a complete knock-out. The *Vsx2^LacZ^* allele exhibits loss-of-function characteristics and adds to the genetic toolkit for understanding *Vsx2* function.

## Introduction

Visual System Homeobox (VSX) genes encode paired-like homeodomain transcription factors required for visual system development in both vertebrates and invertebrates. Two paralogous genes, *Vsx1* and *Vsx2* (formerly *Chx10*), are present in vertebrates and have distinct roles in eye growth and retinal formation during development. In humans, clinical features of *Vsx1* disruption include keratoconus and visual deficits whereas *Vsx2* disruption include non-syndromic congenital bilateral microphthalmia and blindness. *Vsx2* mutations also cause non-syndromic microphthalmia in other vertebrates. Understanding how *Vsx2* functions and is regulated has provided important insights into ocular development.

*Vsx2* expression is activated in the nascent retinal domain during the morphological transition from the optic vesicle to optic cup, remains expressed in retinal progenitor cells (RPCs) through development, and becomes restricted to bipolar cells and Müller glia in the adult retina.^1,2^ Consistent with onset of expression, reductions in eye size and defects in retinal development commence soon after optic cup formation in murine *Vsx2* mutants^1,3^ and at an equivalent stage in *Vsx2* mutant human organoids.^4^ In mice, retinal defects include ectopic expression of non-retinal genes and differentiation programs more akin to neighboring tissues (lineage infidelity), slowed proliferation, delayed onset of neurogenesis, disrupted laminar morphogenesis, and a lack of bipolar cells.^1,3,5–12^

Multiple *Vsx2* alleles encoding structural or regulatory variants exist in mice (informatics.jax.org; Quick Search term: Vsx2; Category: Alleles), but most phenotypic characterizations of *Vsx2* disruption in the eye have been done with three protein coding variants. The most extensively studied is the *ocular retardation J* allele (*orJ;* MGI:1856112), a spontaneous mutation that replaces tyrosine at position 176 with a premature stop codon (Fig. 1A).^1,5^ Although *Vsx2* mRNA remains expressed in the *orJ* mutant retina, VSX2 protein is not detected, and the *orJ* allele is considered a functional null.^1,3,10^ The other two alleles are gene targeted missense mutations that are homologous to human variants associated with congenital microphthalmia and blindness: a glutamine substitution for arginine at amino acid position 200 (*R200Q*; MGI:5449355), and a tryptophan substitution of arginine at position 227 (*R227W*; MGI:5449356; (Fig. 1A)).^3^ Like the *orJ* allele, both are recessive, and their phenotypes are consistent with disrupted *Vsx2* function although they differ in severity. The *R200Q* mutant closely phenocopies *orJ,* while the *R227W* mutant is more severe as revealed by even smaller eyes, increased pigmentation of the retina, and a failure to initiate neurogenesis.^3^ The *R200Q* and *R227W* alleles also differ from *orJ* in that the mutant proteins are expressed.^3^ The *R200Q* mutation is in the homeodomain and abrogates DNA binding to canonical sites.^3,13^ The *R227W* mutation is in the CVC domain, a conserved stretch of approximately 50 amino acids that is a defining feature of the VSX gene family.^3^ Although its mechanism of action remains elusive, the CVC domain also appears to promote DNA binding and sumoylation.^14,15^ Consistent with its role in DNA binding, the *R227W* protein exhibits weakened binding to canonical VSX2 binding sites.^3^

**Figure 1.**
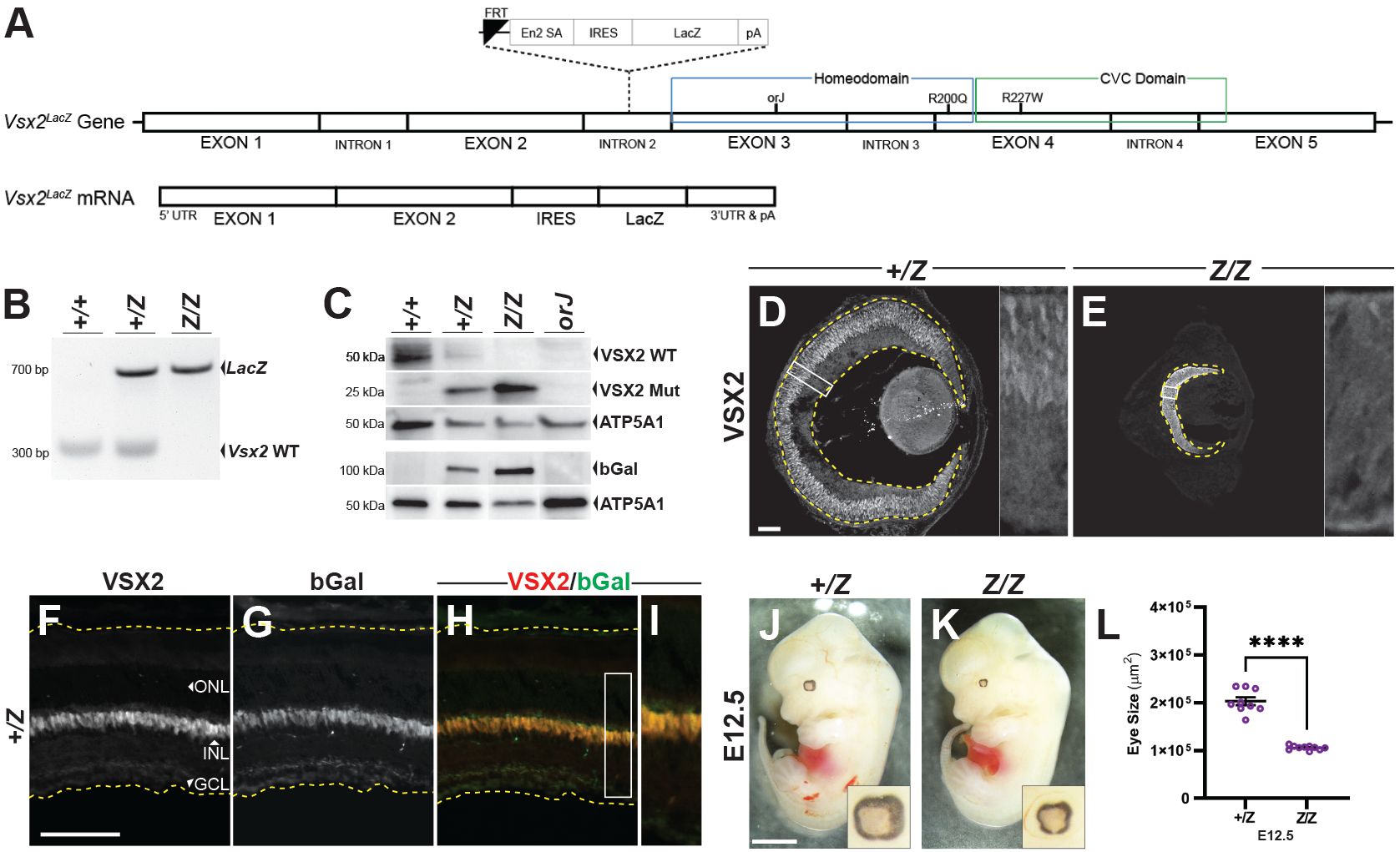
*Vsx2^LacZ^* recombination confirmation and gross phenotype. **(A)** Schematic of *Vsx2^LacZ^* allele and resultant mRNA following Cre recombination. **(B)** Genomic DNA PCR shows mendelian inheritance of *Vsx2* wildtype and *LacZ* sequences confirming germline transmission. **(C)** Western blot analysis of E15.5 retinal lysates. The N-terminal specific VSX2 antibody was used in the top 2 rows to detect full length and truncated VSX2 proteins. B-GAL antibody was used in 4^th^ row to detect b-GAL expression from *LacZ allele*. VSX2 and b-GAL antibodies were used on separate blot. The probing for ATP5A1 serves as a loading control for each blot. **(D-E)** Immunohistology at E15.5 reveals continued yet differentially localized expression of VSX2 in the *LacZ* mutant (retinas are contained within the dashed lines). **(F-I)** Recombination reporter, b-GAL, is co-expressed with VSX2 in the *LacZ* heterozygote INL at P28. **(J-L)** Microphthalmia phenotype of the *LacZ* mutant is observable and significant at E12.5 (****p<0.0001, T-test, sample sizes: 9 (*Z/+*), 10 (*Z/Z*)). Abbreviations: ONL, outer nuclear layer; INL, inner nuclear layer; GCL, ganglion cell layer. Scale bars: (D-H) 100 µm; (J,K) 2000 µm.

Here, we expand the genetic toolkit for *Vsx2* with a new targeted allele that replaces exon 3 with an IRES-*LacZ* expression cassette (*Vsx2^LacZ^*). We show that b-GAL protein expression from the *LacZ* cassette is a reliable reporter of endogenous VSX2 expression, and that the ocular phenotype of the *Vsx2^LacZ^* homozygous mutant (*LacZ* mutant; Z/Z) exhibits the hallmarks of *Vsx2* disruption in a manner similar to the *orJ* mutant. However, unlike *orJ*, the predicted truncated protein is expressed and there are subtle differences in the *LacZ* mutant phenotype – leaving open the possibility that the allele is not a complete knockout. This allele adds to the repertoire of *Vsx2* mutants in which protein variants are expressed and provides an additional resource for probing *Vsx2* function *in vivo*.

## Results

The VSX2-encoding portion of the *Vsx2^LacZ^* mRNA is solely composed of exons 1 and 2, therefore lacking the homeodomain and CVC domain. This is followed by an internal ribosome entry sequence (IRES) and *LacZ* cDNA, which encodes the *Escherichia coli* b-GAL protein (Fig. 1A). The *Vsx2^LacZ^* (*LacZ*) allele was generated by Cre recombination of the *Vsx2^tm1a(EUCOMM)Wtsi^* allele (MGI ID:4453691),^16^ and genomic DNA PCR confirmed germline transmission (Fig. 1B). Western blot analysis of E15.5 retinal lysates probed with an antibody specific to the N-terminal portion of the human VSX2 (aa1-131; see Table 1 for antibodies and dilutions) revealed that like the *orJ* allele, full length VSX2 is not expressed from the *LacZ* allele (Fig. 1C; top row). Unexpectedly, the predicted, truncated VSX2 protein from the *LacZ* allele was detected, indicating stable expression (Fig. 1C; second row). Detection of b-GAL confirmed that the gene targeting strategy was successful and that the IRES was sufficient to produce two distinct proteins from the same transcript (Figure 1C; fourth row). Immunohistology revealed that the mutant protein lacked nuclear localization, consistent with the retention of the nuclear export signal (NES) in exon 2 and the loss of the nuclear localization signal (NLS) in exon 3 (Fig. 1D,E).^17^ We also confirmed that b-GAL accurately reports VSX2 expression, indicated by their extensive colocalization throughout development and easily visualized in the inner nuclear layer (INL) of *LacZ* heterozygous retinas (*LacZ* het; *+/Z*) at P28. (Fig. 1F-I; Supplemental Fig. 1A-F). Finally, the stereotypical microphthalmic phenotype was evident and significant at E12.5 (Fig. 1J-L; see figure legends and Supplemental Table 1 for statistics; see Methods for quantification methods and statistical analyses).

**Table 1:** Primary Antibodies.

| NAME | SPECIES<sup>1</sup> | DILUTION<sup>2</sup> | SOURCE | CATALOG # | RRID |
| --- | --- | --- | --- | --- | --- |
| ATOH7 | Rb | 1:200 (IH) | Novus Biologicals | NBP1-88639 | AB_11034390 |
| b-GAL | Ms | 1:200 (IH)<br>1:1000 (WB) | Developmental Studies<br>Hybridoma Bank | 40-1a-s | AB_528100 |
| BHLHB5 | Rb | 1:200 (IH) | Santa Cruz<br>Biotechnology | sc-6045 | AB_2065343 |
| CCND1 | Rb | 1:400 (IH) | Abcam | ab16663 (SP4) | AB_443423 |
| LHX2 | Rb | 1:300 (IH) | GeneTex | GTX129241 | AB_2783558 |
| MCM6 | Gt | 1:300 (IH) | Santa Cruz<br>Biotechnology | sc-9843 | AB_2142543 |
| MITF | Ms | 1:800 (IH) | Exalpa Biologicals Inc. | X1405M | - |
| OTX1/2 | Gt | 1:1000 (IH) | GeneTex | AF1979 | AB_2157172 |
| PAX2 | Rb | 1:100 (IH) | Covance Laboratories | PRB-276P | AB_291611 |
| PAX6 | Rb | 1:300 (IH) | BioLegend | 901301 | AB_2565003 |
| PCNA | Ms | 1:500 (IH) | Santa Cruz<br>Biotechnology | sc-56 | AB_628110 |
| PKC | Rb | 1:200 (IH) | Sigma-Aldrich | P-4334 | AB_477345 |
| POU4F2 | Gt | 1:200 (IH) | Santa Cruz<br>Biotechnology | sc-6026 | AB_673441 |
| RECOVERIN | Rb | 1:100 (IH) | Chemicon International | AB5585 | AB_2253622 |
| SOX2 | Rb | 1:400 (IH) | Abcam | ab97959 | AB_2341193 |
| SOX9 | Rb | 1:1000 (IH) | EMD Serono Inc | AB5535 | AB_2239761 |
| TUBB3 | Rb | 1:40K (IH) | BioLegend | 802001 | AB_2564645 |
| VSX1 | Rb | 1:250 (IH) | Levine Lab<sup>26</sup> | - | - |
| VSX2 - NT | Sh | 1:400 (IH)<br>1:1000 (WB) | Exalpa Biologicals Inc. | X1180P | AB_2314191 |
| ATP5A1 | Rb | 1:2000 (WB) | Proteintech | 14676-1-AP |  |
| <sup>1</sup> Gt, goat; Ms, mouse; Rb, rabbit; Sh, sheep |  |  |  |  |  |
| <sup>2</sup> IH, immunohistology; WB, western blot |  |  |  |  |  |

We next performed immunohistology to characterize the *LacZ* mutant phenotype in the retina. Establishment of optic cup regionalization appeared intact at E10.5 as evidenced by VSX2 expression in the retina, LHX2 and PAX6 in the RPE and retina, OTX1/2 and MITF in the RPE (Fig. 2A-E,G-K). The RPC and proliferation markers PCNA and CCND1 also appeared to be normal in the E10.5 *LacZ* mutant retina (Supplemental Fig. 2A-D). However, MITF was ectopically expressed in the retina (Fig. 2E,K) and PAX2 expression extended into the dorsal retina in the *LacZ* mutant (Fig. 2F,L; arrows). At E11.5, VSX2, LHX2, and PAX6 expression remained unaltered (Fig. 2M-O,S-U), but the ectopic expression of MITF (Fig. 2E,K) and dorsal expression of Pax2 (Fig. 2R,X; arrows) were still evident, and OTX1/2 expression in the central retina was elevated in the *LacZ* mutant (Fig. 2P,V). These altered expression patterns are consistent with compromised retinal identity. Further evidence of altered retinal identity was observed at E12.5 with continued ectopic expression of OTX1/2 in the peripheral retina, most notably on the ventral side (Fig. 4E,K; arrows). From E11.5 to E15.5, the RPC population, as indicated by PCNA, is maintained (Fig. 3A-F; Supplemental Fig. 2E,F), but proliferation progressively decreased, most notably in the peripheral retina, as assessed by reduced CCND1 expression (Fig. 3G-L; Supplemental Fig. 2G,H; arrows) and EdU incorporation (Fig. 3M-R; Supplemental Fig. 2I,J; arrows). This early embryonic phenotype emulates those of the *orJ*, *R200Q,* and *R227W* mutants.^1,3^

**Figure 2.**
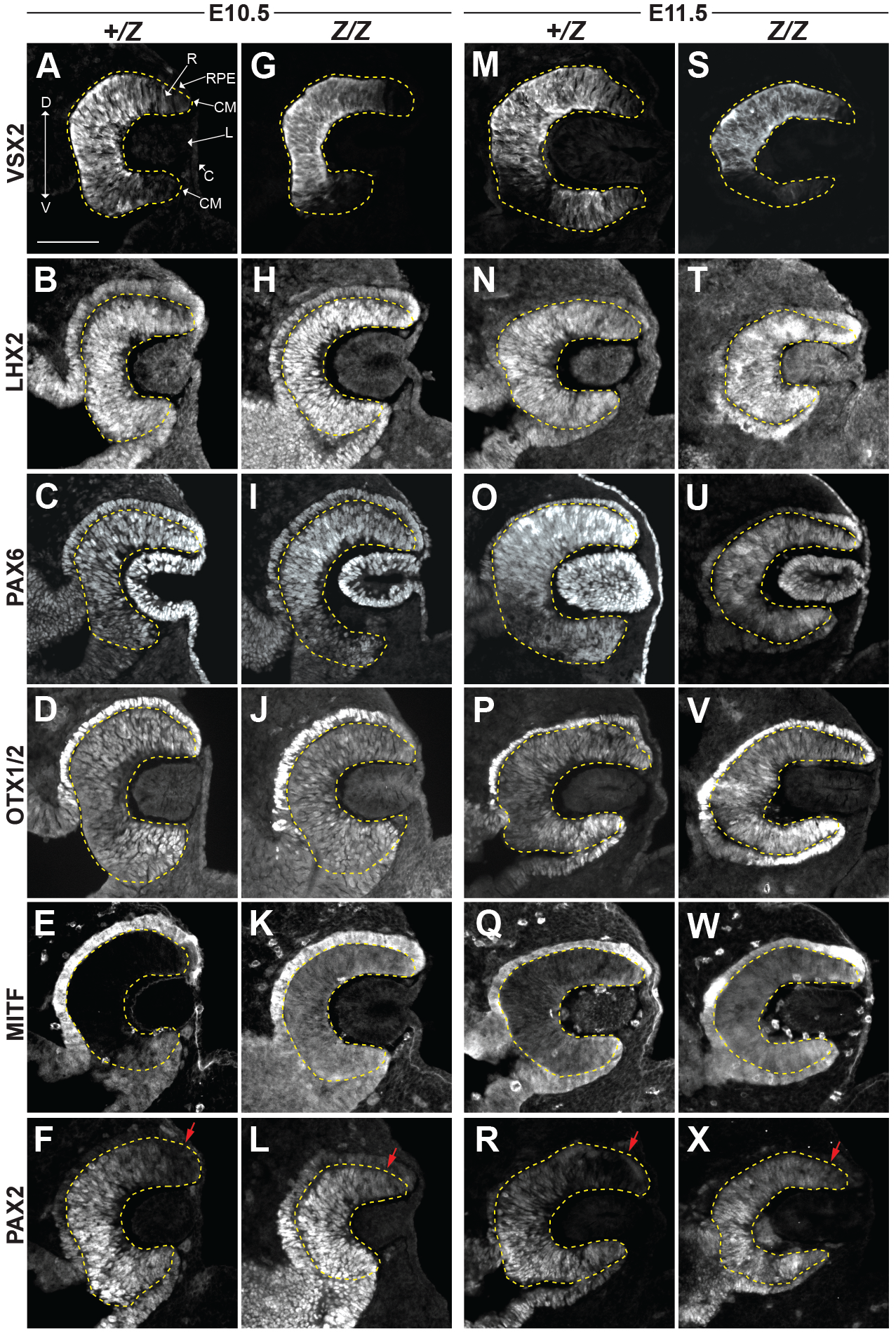
Regionalization of the early embryonic *LacZ* mutant eye. **(A-L)** Markers of eye field specification and regionalization at E10.5. All markers exhibit similar expression patterns in the *LacZ* mutant compared to the *LacZ* het except for MITF, which is ectopically expressed in the retina (E,K) and PAX2, which is expanded into the dorsal retina (arrows; F,L). **(M-X)** Markers of eye field specification and regionalization at E11.5. MITF continues to be ectopically expressed (Q,W) and PAX2 extends into the dorsal retina of the *LacZ* mutants (arrows; R,X). All other markers were similar between the *LacZ* het and mutant. Abbreviations: R, retina; RPE, retinal pigment epithelium; CM, ciliary margin; L, lens; C, cornea. Scale bar: 100 µm.

**Figure 3.**
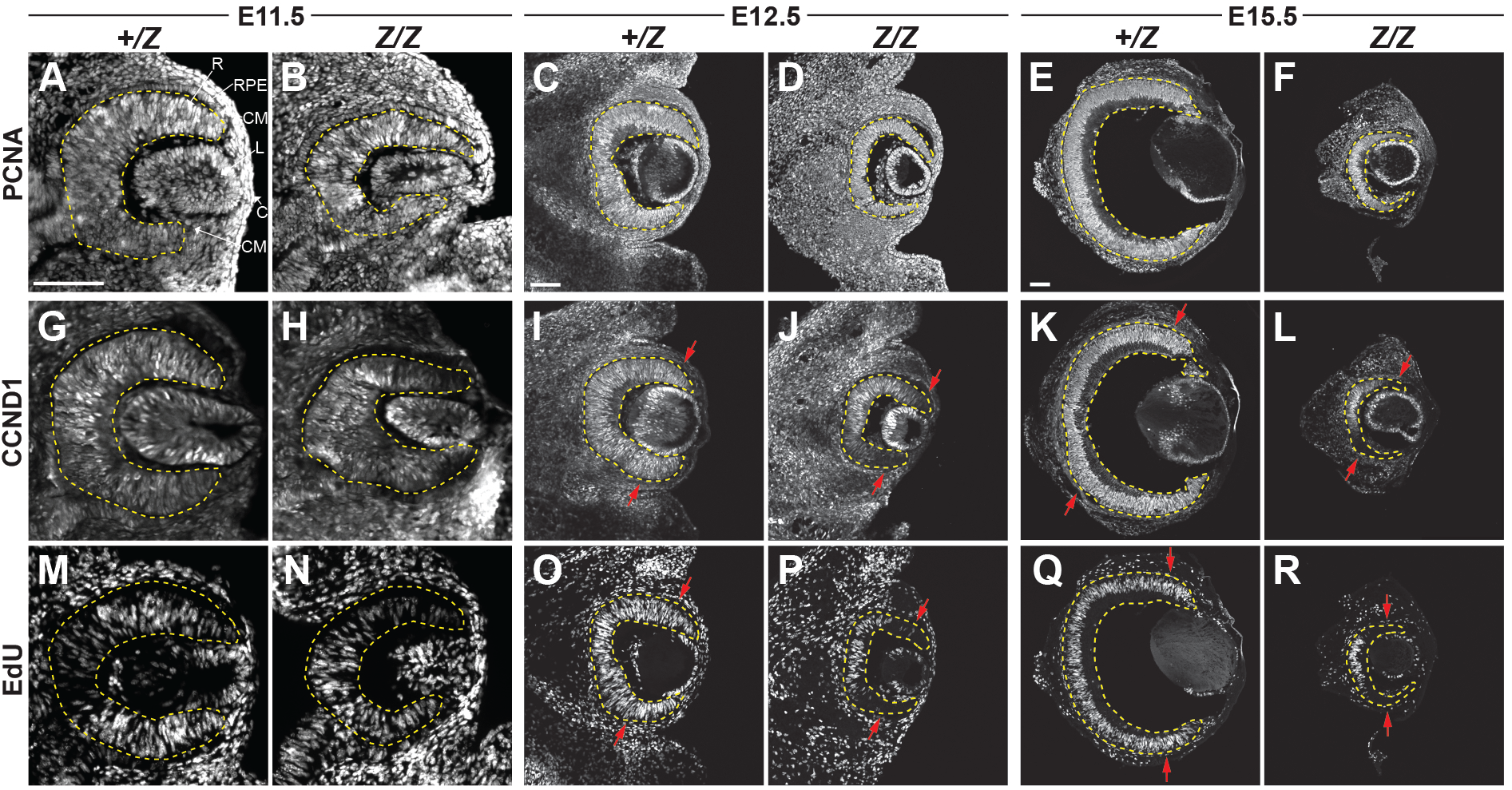
Proliferation is reduced in the embryonic *LacZ* mutant retina. **(A-F)** The general proliferation marker PCNA remains expressed throughout the *LacZ* mutant retina, similar to the *LacZ* het at E11.5, E12.5, and E15.5. In contrast, the patterns of CCND1 expression **(G-L)** and EdU incorporation **(M-R)** became progressively restricted to the central retina in the *LacZ* mutants compared to *LacZ* hets, most notably at E12.5 and E15.5. Arrows in I-L and O-R point to the regions where staining is absent in the *LacZ* mutant retina compared to the *LacZ* het. Scale bar: 100 µm.

**Figure 4.**
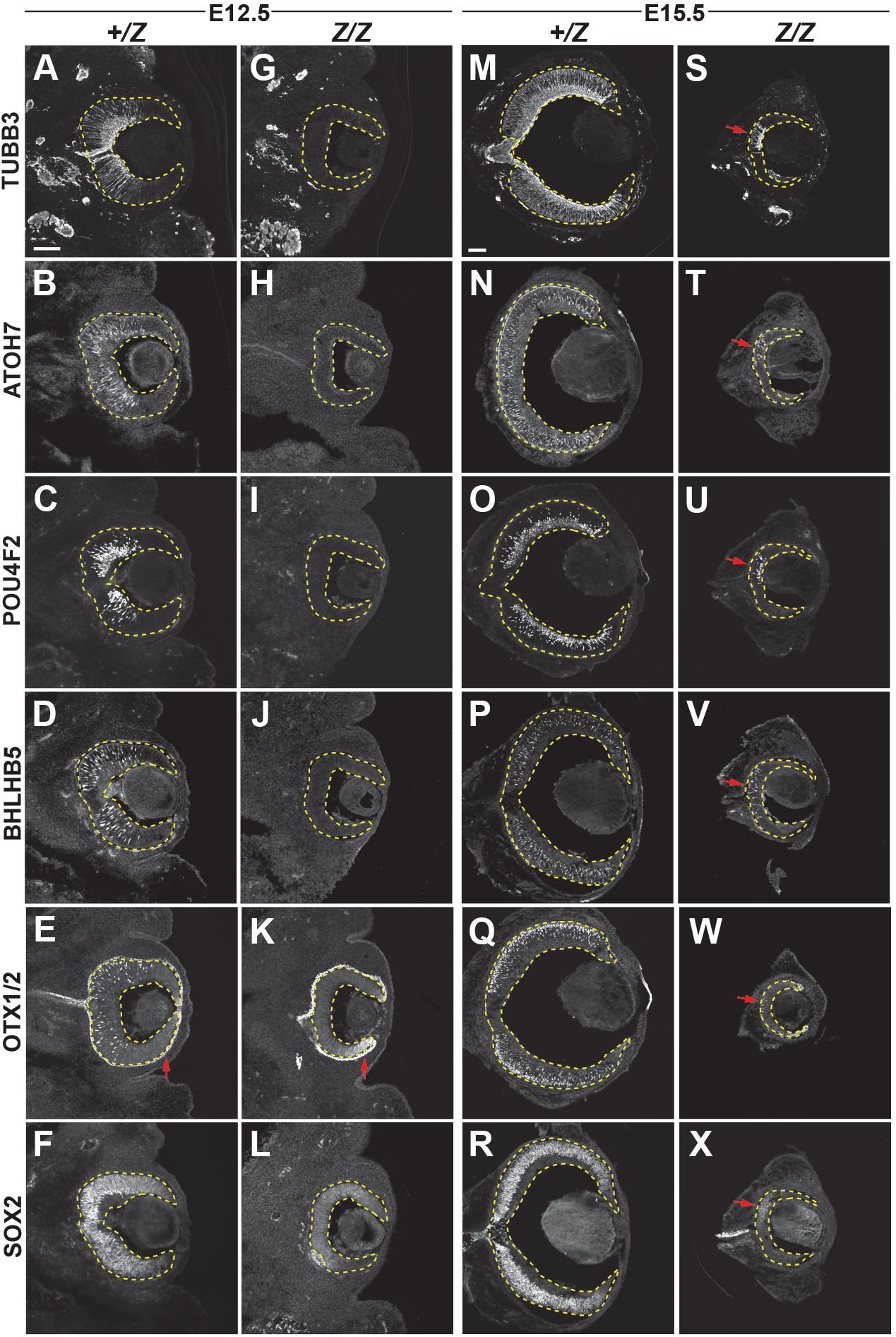
Retinal neurogenesis is delayed and centrally restricted in the *LacZ* mutant. Immunohistology assessing cytoskeletal maturation with TUBB3 **(A,G,M,S)**, the presence of neurogenic progenitors with ATOH7 **(B,H,N,T)**, retinal ganglion cells with POU4F2 **(C,I,O,U)**, amacrine cells with BHLHB5 **(D,J,P,V)** and photoreceptor precursors with OTX1/2 (**E,K,Q,W)**, reveal a delay in neurogenesis in the *LacZ* mutant retina, with markers lacking at E12.5 and centrally restricted at E15.5 (arrows in S-W). OTX1/2 staining also reveals ectopic staining in the ventral peripheral retina at E12.5 (compare red arrows in E and K), which is associated with compromised retinal identity. **(F,L,R,X)** Expression of neurogenic competence regulator SOX2 in RPCs is reduced at E12.5 (F,L) and centrally restricted at E15.5 in the *LacZ* mutant retina (R,X; arrow denotes region of persistent SOX2 expression). Scale bar: 100 µm.

Neurogenesis and establishment of the inner and outer neuroblastic layers initiate at E11.5 in the central retina and progress out to the retinal peripheral margin by E15.5. This is readily observed in the *LacZ* het retina at E12.5, E13.5, and E15.5 using the expression of neurogenesis and cell-type specific markers including TUBB3, a marker of cytoskeletal maturation in early post mitotic neuronal precursors, ATOH7, a marker expressed by neurogenic progenitors preceding cell cycle exit, BHLHB5 and POU4F2, markers of amacrine and RGCs respectively, and OTX1/2, a marker of neurogenic RPCs biased towards photoreceptor fates as well as photoreceptor precursors (Fig. 4A-E;M-Q; Supplemental Fig. 2K,M,O,Q,S). In contrast, none of these neurogenesis markers were detected in the central retina of the *LacZ* mutant at E12.5 (Fig. 4G-K). OTX1/2 was expressed in the peripheral ventral retina (Fig. 4E,K; arrows), but as mentioned above, this expression is associated with compromised retinal identity.^7^ By E15.5 in the *LacZ* mutant, all markers were expressed in patterns consistent with neurogenesis, although stained cells were restricted to the central retina (Fig. 4S-W; arrows). Correlated with the delayed neurogenesis is reduced expression of the neurogenic competence marker SOX2 at E12.5 (Fig. 4F,L), and restricted expression in a small central region of the *LacZ* mutant retina at E13.5 (Supplemental Fig.2 S,T; arrows) and E15.5 (Fig. 4R,X; arrows).^17^ These observations are consistent with a delay in the onset of retinal neurogenesis, to a similar extent as the *orJ* and *R200Q* mutants.

We observed variation in the severity of the microphthalmic phenotype of *LacZ* mutant embryos at E15.5, ranging from similar to more severe than in *orJ* embryos (Fig. 5A). Although the mean reduction in eye size of the *LacZ* and *orJ* mutants were similar in magnitude when compared to their respective controls (Fig. 5B, Supplemental Table 1), variability was greater in the *LacZ* mutants as determined by the coefficients of variation (CV) for each genotype (Fig. 5C; see Methods for formula). We also observed variation in the extent of neurogenesis, as assessed by TUBB3 expression, between *LacZ* mutants (Fig. 5D-G). While neurogenic length and density (defined in methods) were similar in the *LacZ* and *orJ* mutants (Fig. 5H,I; Supplemental Table 1), the extent of variation in these measurements were similar except for neurogenic length in the *LacZ* mutants, which had a larger CV compared to the other genotypes (Fig. 5J,K). Linear regression analysis revealed that neurogenic density and eye size were highly correlated (Fig. 5L), but this was not the case for neurogenic length and eye size (Fig. 5M). These observations suggest a link between eye size and neuron production, but not between eye size and progression of the neurogenic wave from the central to peripheral retina.

**Figure 5.**
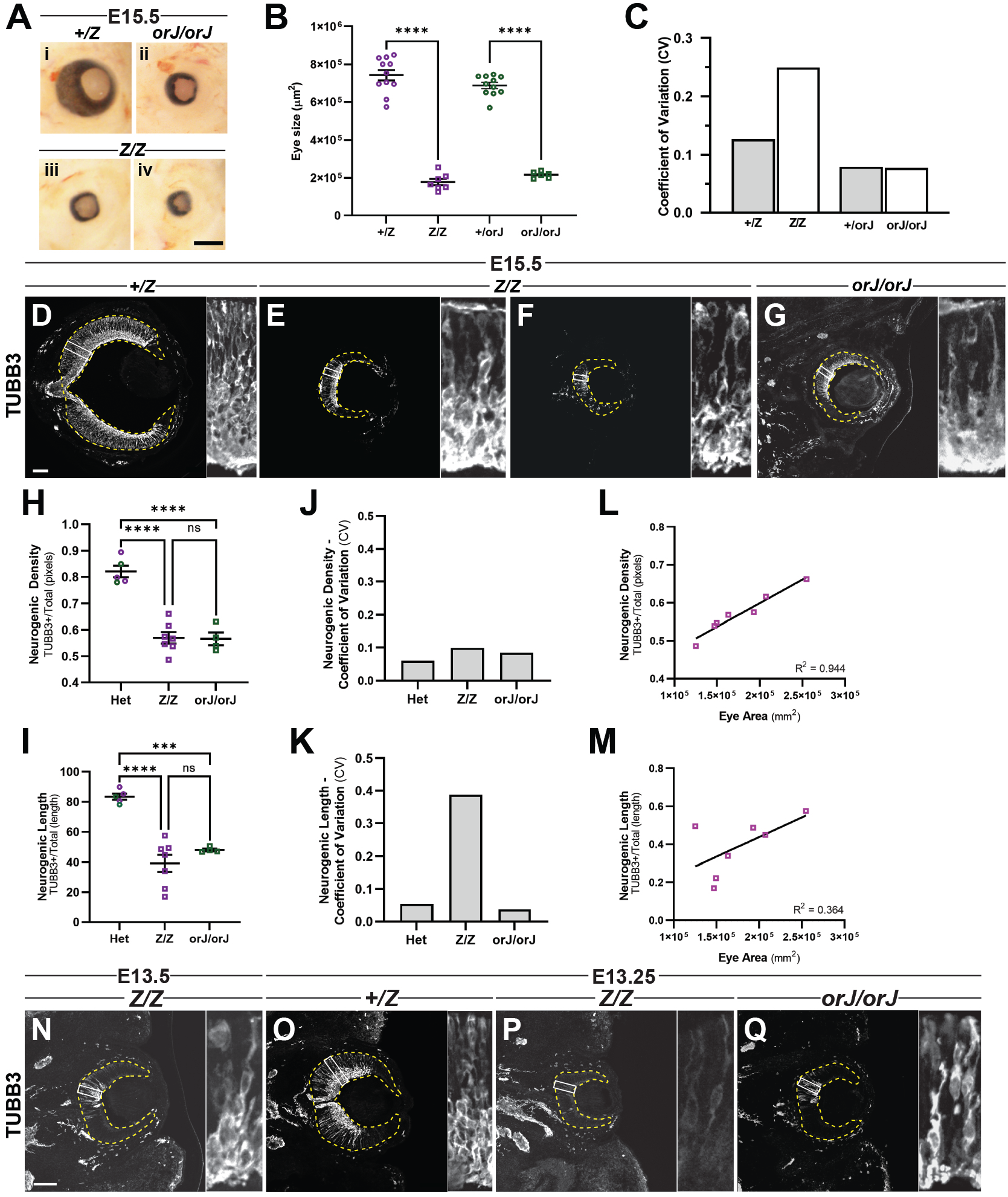
Timing of neurogenesis initiation and E15.5 variation in *LacZ* mutant deviate from the *orJ* mutant phenotype. **(A)** Microphthalmia was similar in the *LacZ* and *orJ* mutants at E15.5, but severity was variable in the *LacZ* mutants (compare iii and iv). **(B,C)**. Quantifications of eye size revealed similar degrees of severity (B; ****p<0.0001; ANOVA; sample sizes: 11 (*Z/+*), 7 (*Z/Z*), 11 (*orJ/+*), 6 (*orJ/orJ*)), but the coefficient of variation (CV) was highest in the *LacZ* mutants compared to the other genotypes (C). **(D-G)** TUBB3 expression, indicating neurogenesis, is centrally restricted and seemingly covaries with microphthalmia severity in the *LacZ* mutant at E15.5. **(H,I)** Neurogenic density (H); and length (I) are similarly decreased in the *LacZ* and *orJ* mutants (****p<0.0001; ***p<0.001; ^ns^p>0.05; ANOVA; sample sizes: 5 (*Z/+* and *orj/+* combined (Het)), 7 (*Z/Z*), 4 (*orJ/orJ*)). **(J,K)**. CVs for neurogenic density were similar across genotypes (J), but the CV for neurogenic length was specifically increased in the *LacZ* mutant (K). **(L)** Linear regression analysis of eye size and neurogenic density in *LacZ* mutant retinas at E15.5 (R^2^=0.944; F=83.44; p=0.0003; sample size: 7). **(M)** Linear regression analysis of eye size and neurogenic length in *LacZ* mutant retinas at E15.5 (R^2^=0.364; F=2.856; p=0.1518; sample size: 7). **(N-Q)** Neurogenesis initiates at E13.5 in *LacZ* mutant (compare N to P). In contrast, neurogenesis initiates in the *orJ* retina by E13.25 (Q). Scale bars: (A) 500 µm; (D-G; N-Q) 100 µm.

Given the variation in neurogenesis at E15.5, we hypothesized that differences between the *LacZ* and *orJ* mutant phenotypes began earlier. We therefore focused on the onset of neurogenesis, which exhibits a near stereotypical delay of two days in the *orJ* retina, initiating at approximately E13.5.^8^ Expression of TUBB3 (Fig. 5N; box and inset) and OTX1/2 (Supplemental Fig. 2R; arrow) was observed at E13.5 in the central retina of the *LacZ* mutant, but cells were sparse compared to the *LacZ* het (Supplemental Fig. 2Q). Similarly, cells expressing the neurogenic progenitor marker ATOH7 were present but sparse in the *LacZ* mutant retina (Supplemental Fig. 2K,L; arrow), whereas POU4F2, and BHLHB5 were not detected (Supplemental Fig. 2M-P). We therefore examined TUBB3 staining in retinas from the *LacZ* and *orJ* mutants at E13.25. Indeed, neurogenesis was not observed at E13.25 in the *LacZ* mutant retina (n=7) but was in the *orJ* mutant retina (n=4; Fig. 5O-Q). The sum of these observations suggests that the *LacZ* mutant exhibits subtle increases in phenotypic severity compared to the *orJ* mutant (see discussion).

Elevated MITF expression in the retina is a central driver of the ocular phenotypes in *orJ, R200Q,* and *R227W* mutant mice, and a semi-dominant negative allele of *Mitf* (*Mitf^mi^*) is sufficient to significantly reduce phenotypic severity.^3,8,9,18^ We predicted a similar outcome with the *LacZ* allele, and to test this, we crossed *LacZ* mutant mice with our *Vsx2^orJ/orJ^*; *MITF^mi/+^* line to generate *Vsx2^orJ/Z^*; *MITF^mi/+^* embryos. At both E12.5 and E15.5, *Vsx2^orJ/Z^*; *MITF^mi/+^* eyes were significantly larger than those of *Vsx2^orJ/Z^*; *MITF^+/+^* (Fig. 6). This suggests that, like the other *Vsx2* mutants, *Mitf* has a prominent role in causing microphthalmia in the *LacZ* mutant retina.

**Figure 6.**
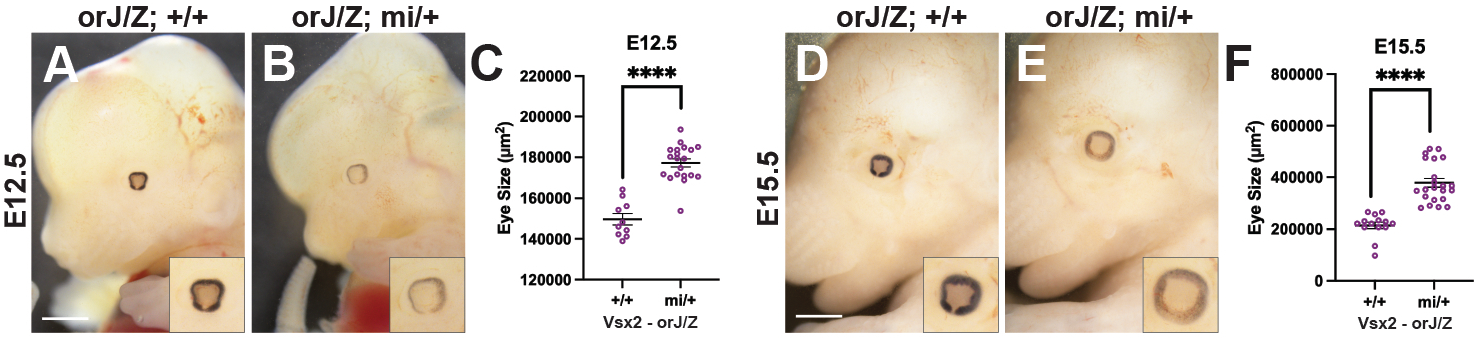
Functional rescue of embryonic *LacZ* mutant microphthalmic phenotype with *Mitf^mi/+^* allele. The semi-dominant negative allele *MITF^mi/+^* reduced the severity of microphthalmia in *Vsx2^orJ//Z^* mutants at E12.5 **(A-C)** and E15.5 **(D-F)**. ****p<0.0001; T-test; sample sizes for E12.5: 10 (*orJ/Z;+/+*),20 (*orJ/Z;mi/+*); sample sizes for E15.5: 14 (*orJ/Z;+/+*), 22 (*orJ/Z;mi/+*)). Scale bar: 1000 µm.

**Table 2:** Secondary Antibodies.

| NAME | DILUTION<sup>1</sup> | SOURCE | CATALOG # | RRID |
| --- | --- | --- | --- | --- |
| Alexa Fluor™ 488 donkey anti-rabbit IgG (H+L) | 1:1000 (IH) | Invitrogen | A21206 | AB_2535792 |
| Alexa Fluor™ 568 donkey anti-rabbit IgG (H+L) | 1:1000 (IH) | Invitrogen | A10042 | AB_2534017 |
| Alexa Fluor™ 647 donkey anti-rabbit IgG (H+L) | 1:1000 (IH) | Invitrogen | A31573 | AB_2536183 |
| Alexa Fluor® 488 Affinipure Donkey Anti-Mouse IgG (H+L) | 1:1000 (IH) | Jackson ImmunoResearch<br>Laboratories Inc. | 715-545-150 | AB_2340846 |
| Alexa Fluor® 594 Affinipure Donkey Anti-Mouse IgG (H+L) | 1:1000 (IH) | Jackson ImmunoResearch Laboratories Inc. | 715-585-150 | AB_2340854 |
| Alexa Fluor® 647 Affinipure Donkey Anti-Mouse IgG (H+L) | 1:1000 (IH) | Jackson ImmunoResearch Laboratories Inc. | 715-605-150 | AB_2340862 |
| Alexa Fluor® 488 Affinipure Donkey Anti-Goat IgG (H+L) | 1:1000 (IH) | Jackson ImmunoResearch Laboratories Inc. | 705-545-147 | AB_2336933 |
| Alexa Fluor® 594 Affinipure Donkey Anti-Goat IgG (H+L) | 1:1000 (IH) | Jackson ImmunoResearch Laboratories Inc. | 705-585-147 | AB_2340433 |
| Alexa Fluor® 647 Affinipure Donkey Anti-Goat IgG (H+L) | 1:1000 (IH) | Jackson ImmunoResearch Laboratories Inc. | 705-605-147 | AB_2340437 |
| Peroxidase AffiniPure™ Donkey Anti-Goat IgG (H+L) | 1:1000 (WB) | Jackson ImmunoResearch Laboratories Inc. | 705-035-147 | AB_2313587 |
| Anti-Rabbit IgG (H+L), HRP Conjugate | 1:1000 (WB) | Promega | W4011 | AB_430833 |
| Anti-Mouse IgG (H+L), HRP Conjugate | 1:1000 (WB) | Promega | W4021 | AB_430834 |
| <sup>1</sup> IH, immunohistology; WB, western blot |  |  |  |  |

The effects of early *Vsx2* dysfunction are compounded in the postnatal retina as seen by severe hypocellularity and tissue disorganization. Additionally, due to the specific postnatal requirement of *Vsx2* for the specification and differentiation of the last neuronal cell class, bipolar cells, they are not generated.^1,6,11,19^ These features are also evident in the *LacZ* mutant retina. At P0 (Fig. 7), the *LacZ* mutant retina is reduced to a 1-3 cell-thick, epithelium consisting of dispersed RPCs and/or newly forming Müller glia (EdU, MCM6, SOX9) and differentiating retinal neurons (TUBB3) including RGC’s (POU4F2), amacrine cells (BHLHB5), and photoreceptors (RECOVERIN, OTX1/2). At P10 (Fig. 8), the thin apical epithelium, composed largely of photoreceptors (RECOVERIN, OTX1/2) remains intact while an additional population of disorganized aberrantly proliferating RPCs or Müller glia (EdU, MCM6, Sox9), and neurons (TUBB3) including RGCs (POU4F2), and amacrine cells (BHLHB5) arises basal to the organized epithelium. Signs of degeneration emerge at P14 (Supplemental Fig. 3), as the cellular organization at the apical surface becomes disrupted and hypocellular regions, possibly plexiform outgrowths, arise in the disorganized basal cell layer. By P28 (Fig. 9), the retina has severely degenerated -consisting of disorganized retinal progenitor cells, and specified Müller glia, RGCs, amacrine cells, and photoreceptors. Consistent with the other *Vsx2* mutants, bipolar cells are not observed in the *LacZ* mutant retina at P14 (Supplemental Fig. 3Q-T) or P28 (Fig. 10) as revealed by PKCα (PKC) and VSX1. Interestingly, sparse PKC-positive cells were observed at P14, which could suggest limited bipolar cell formation, but this antibody also detects amacrine cells and likely accounts for the pattern observed at P28.^6^

**Figure 7.**
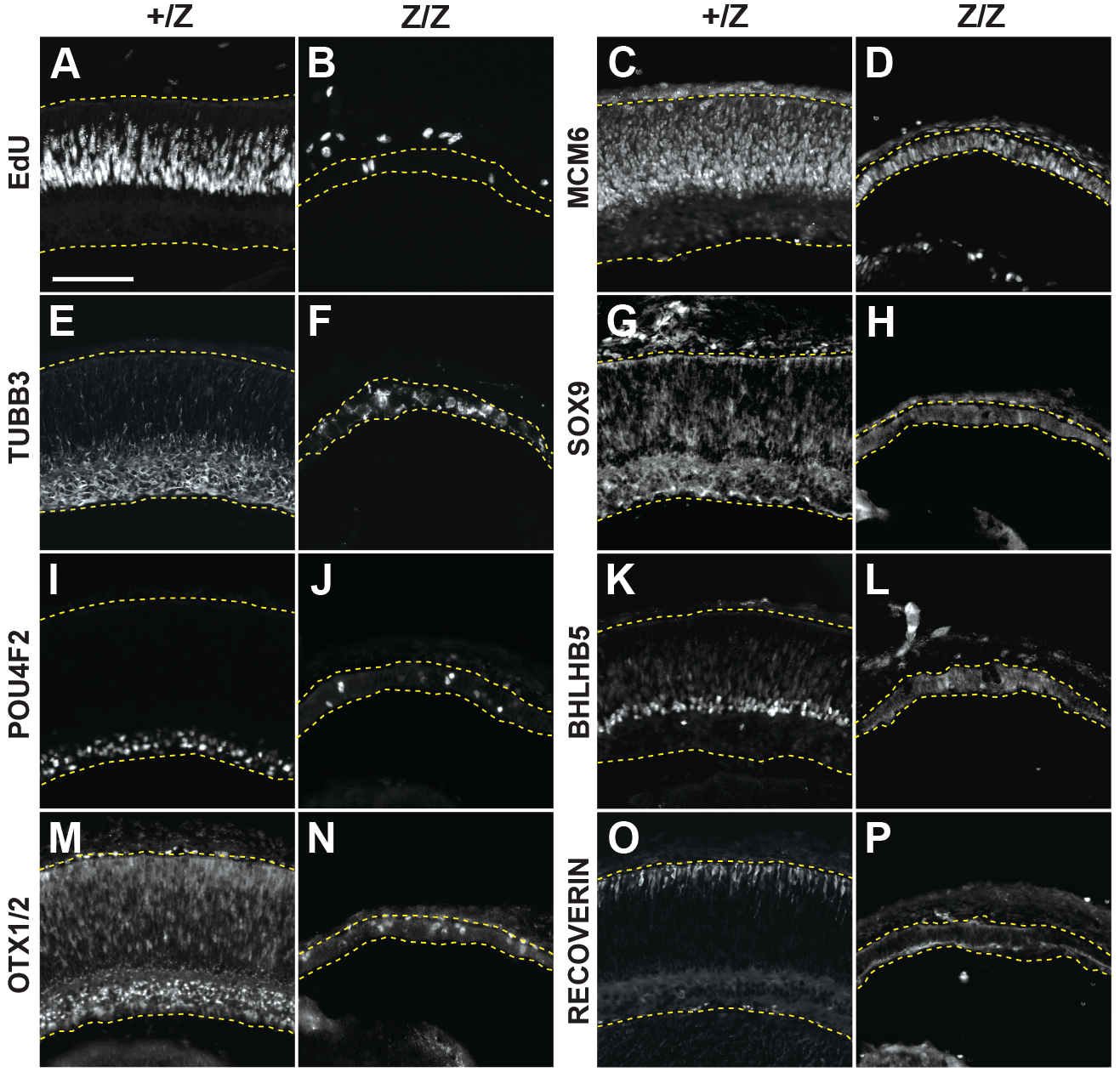
Immunohistology at P0. The *LacZ* mutant retina at P0 is reduced to a thin epithelium, containing interspersed **(A-D,G,H)** proliferating RPCs (EdU, MCM6, SOX9). **(E-P)** Similarly, markers of differentiating cells (TUBB3, POU4F2, BHLHB5, OTX1/2, RECOVERIN are expressed in a disorganized manner. In addition to marking RPCs, SOX9 could indicate precocious Müller glia differentiation. Scale bar: 100 µm.

**Figure 8.**
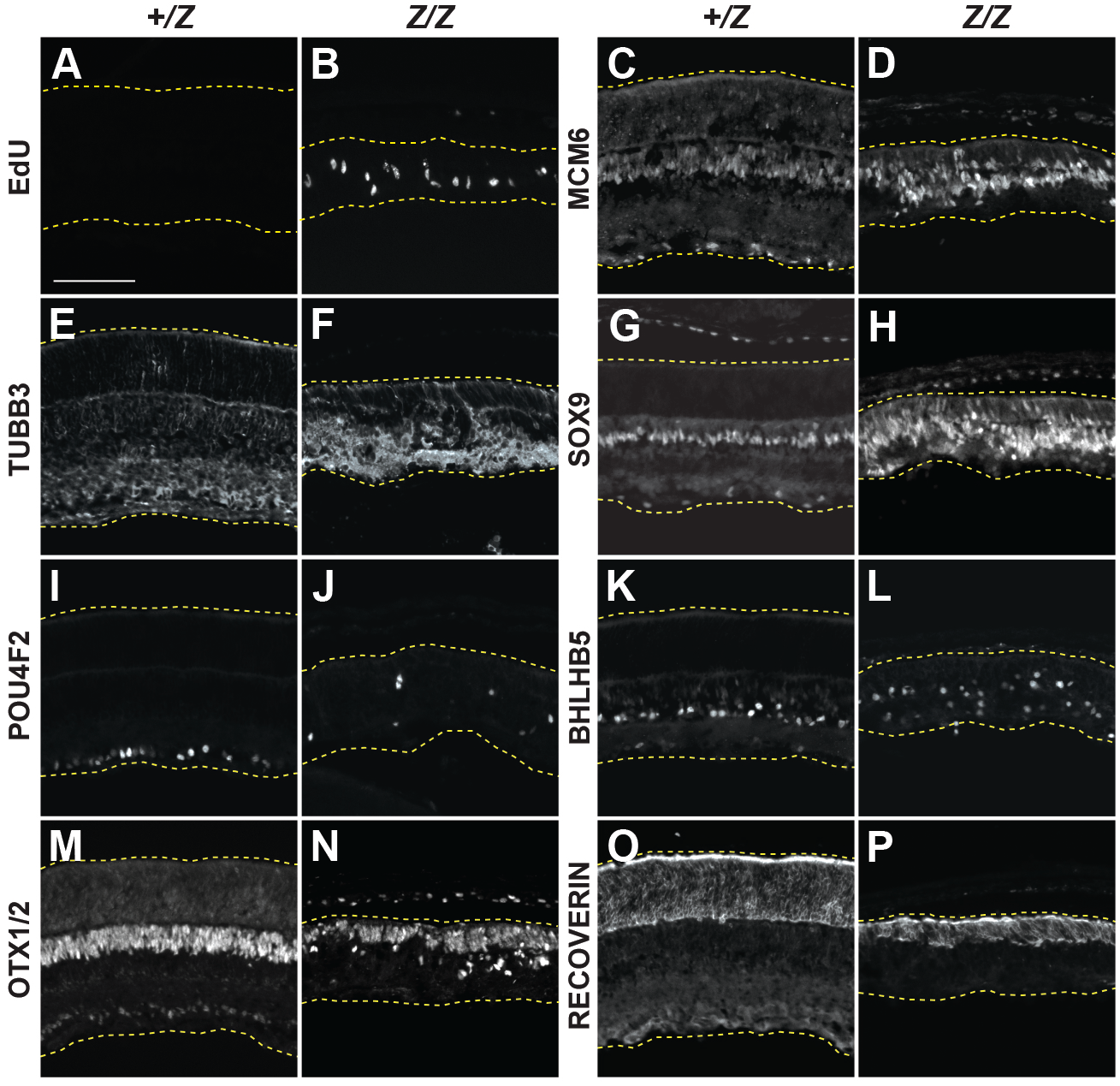
Immunohistology at P10. **(A-L)** A disorganized population of proliferating cells (EdU, MCM6), new neurons (TUBB3, POU4F2, BHLHB5), and Müller glia (SOX9), persist in the *LacZ* mutant retina. (**M-P)** In contrast, the photoreceptor markers OTX1/2 and RECOVERIN suggest the presence of an organized apical epithelium. Scale bar: 100 µm.

**Figure 9.**
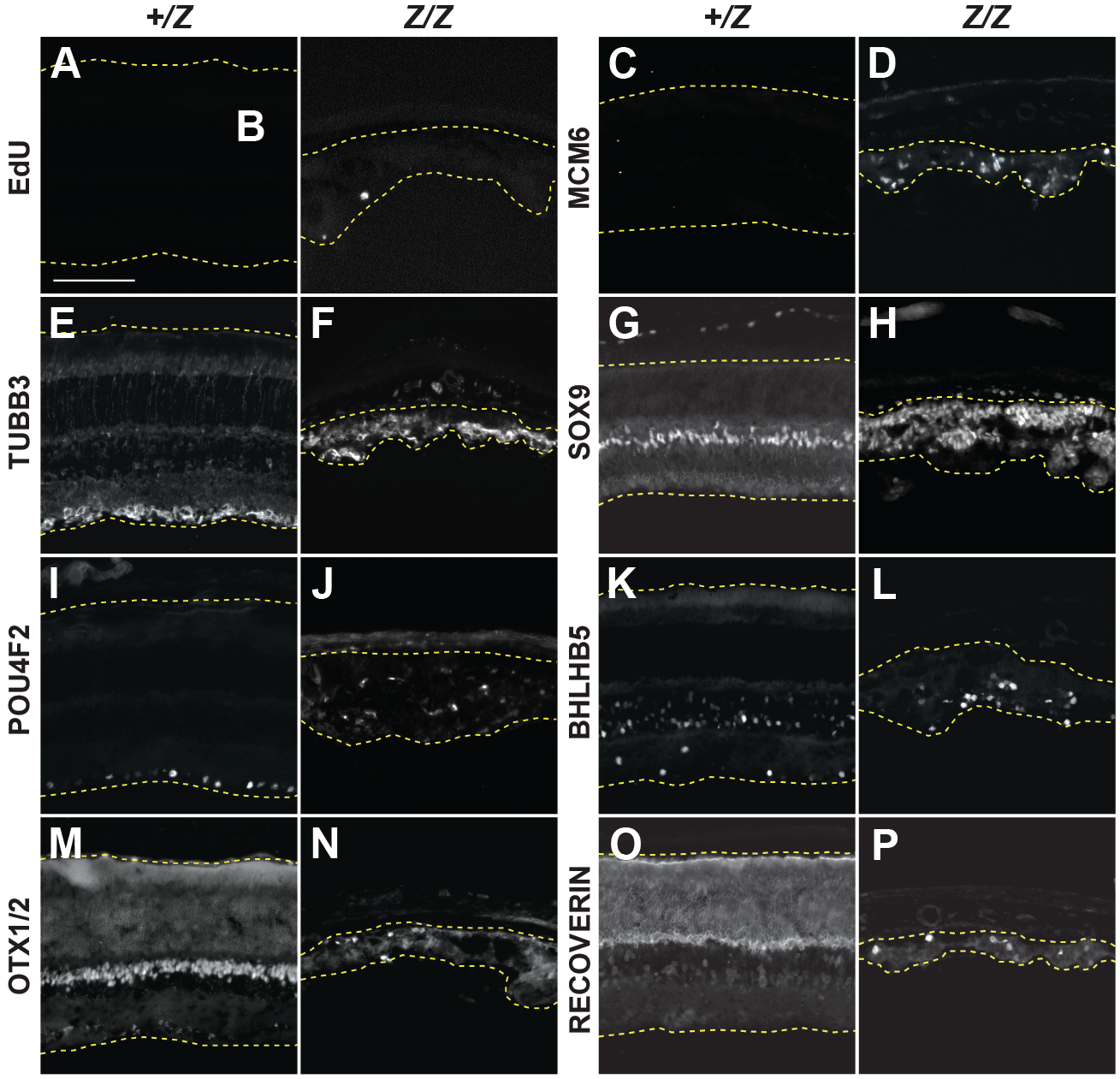
Immunohistology at P28. The *LacZ* mutant retina at P28 is severely degenerated and disorganized. Lacking cellular organization, spherical **(A-D)** aberrantly proliferating cells are interspersed with **(E-P)** progenitors (MCM6, SOX9), differentiating neuronal cells (TUBB3), Müller glia (SOX9), retinal ganglion cells (POU4F2), amacrine cells (BHLHB5), and photoreceptors (OTX1/2, RECOVERIN). Scale bar: 100 µm.

**Figure 10.**
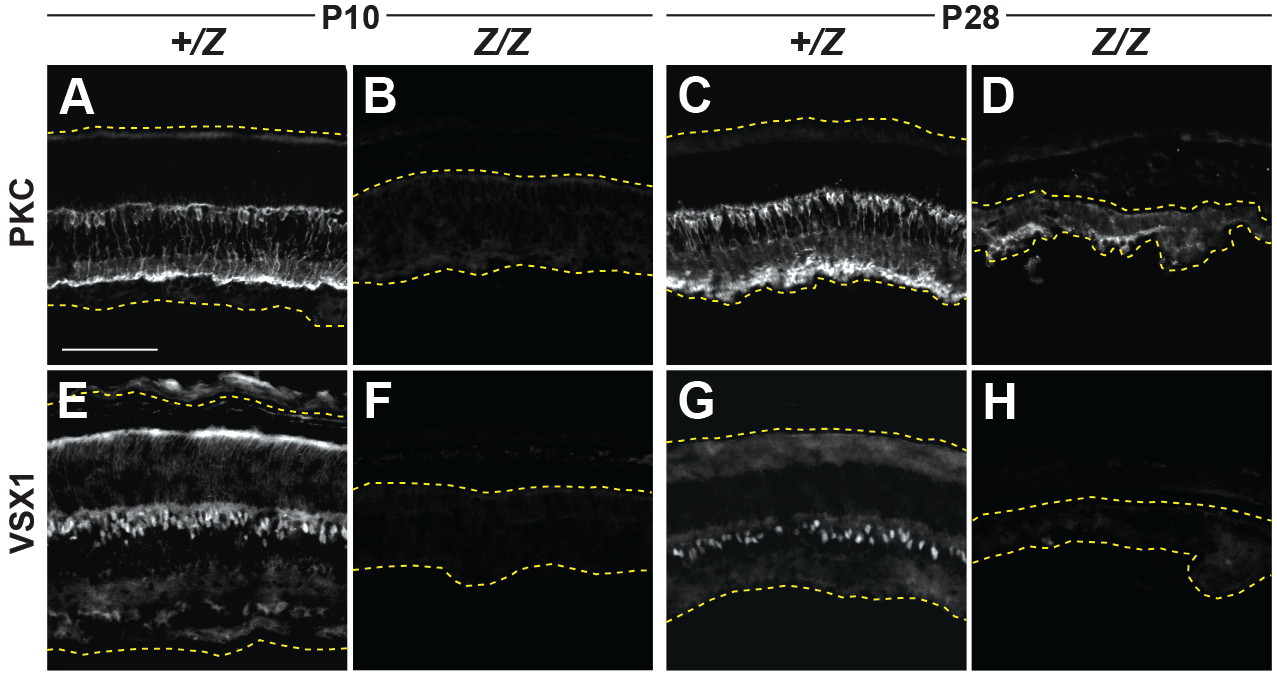
Bipolar cells are not detected in the *LacZ* mutant retina. **(A-D)** The rod-bipolar marker PKC is not detected in the *LacZ* mutant retina at P10, but basally displaced cells are observed at P28. This antibody also detects amacrine cells at this age. **(E,F)** The cone-bipolar marker VSX1 is not detected in the *LacZ* mutant retina at P10 or P28. Scale bar: 100 µm.

## Discussion

Based on the observations reported here, we propose that the *LacZ* allele exhibits loss-of-function properties with strong phenotypic similarities to the *orJ* null allele, which lacks protein expression, and the *R200Q* allele, which produces a DNA binding-deficient protein.^3^ All three mutants exhibit bilateral microphthalmia, reduced retinal proliferation, delayed neurogenesis, an absence of bipolar cells, and are affected by ectopic Mitf activity.^3,8,9,12,18^ Because the mutant VSX2 protein is expressed, we cannot conclude that the *LacZ* allele is a complete knockout, and consistent with this, we observed subtle differences in the timing of neurogenesis onset in the *LacZ* mutant retinas compared to *orJ* mutant, and greater variability amongst *LacZ* mutants at E15.5 when compared to that of *orJ* mutants in both eye size and parameters of neurogenesis.

There are two reasons that could account for these differences. One is that the mutant protein is not inert (see below). The other is that the *LacZ* mutant phenotype is sensitive to its genetic background, which has been documented for the *orJ* mutant phenotype. The *orJ* mutation was discovered in the 129svimj background, but when outcrossed, phenotypic variations occur.^5,11,20^ For example, increased pigmentation and more severe microphthalmia resulted when *orJ* mice were bred with mice with mixed genetic backgrounds including C57Bl6 mice.^21^ We also observed similar increases in phenotypic severity in an outcross with Black Swiss mice (unpublished). In contrast, less severe phenotypes were observed when *orJ* mice were crossed to *p27Kip1* (*Cdkn1b*) knockout mice on a C57BL/6 genetic background^6^ or outcrossed with wild type *Mus castaneous*.^11^ Relevant here is that our characterization of the *LacZ* allele was done in a mixed C57BL/6;129sv mixed genetic background which could contribute to variations in phenotypic severity.

It is also possible that the mutant VSX2 protein has activity, but what this activity might be is not clear. The mutant coding sequence extends from amino acid 1 through 151 (out of 361 or 380; https://www.ncbi.nlm.nih.gov/protein/NP_031727) and lacks the NLS, homeodomain and CVC domain, making direct DNA binding unlikely. The NES is retained, however, and could contribute to non-nuclear localization.^17^ Polyubiquitination and subsequent degradation of zebrafish Vsx1 protein is enhanced by the CVC domain and a conserved, putative PEST sequence located close to the CVC domain, and their absence could explain the apparent stability of the mutant Vsx2 protein.^22^ Interactions of the mutant protein with other proteins is possible, but domains or motifs have not been identified in the retained N-terminal portion of VSX2 other than putative disordered regions. It is also possible that the mutant protein has novel or neomorphic activity because it is no longer localized to the nucleus, but how this might alter the phenotype is hard to reconcile because the allele is not dominant. Interestingly, the *R227W* allele exhibits properties of a recessive neomorph.^3^ We attributed this to the ability of the *R227W* mutant protein, retaining weak DNA binding, to activate a cryptic transcriptional circuit that increased non-retinal gene expression, including *Mitf,* beyond that observed in the *orJ* mutant retina.^3^ Further work is needed to determine if the truncated VSX2 protein from the *LacZ* allele has activity.

### Limitations of the study

It is unclear whether the subtle differences in the phenotypes between the *LacZ* and *orJ* mutants are due to the expression of the mutant protein in the *LacZ* mutant, differences in genetic background, or both. One approach to address this issue is to minimize differences in genetic background by generating a congenic 129sv/imj strain carrying the *LacZ* allele. It’s also unclear if the mutant protein has activity or whether a similar truncated protein with pathogenicity could exist in humans. This limitation could be addressed by generating CRISPR mutations in human iPSCs that are differentiated into retinal organoids. Finally, although eye size is improved in the combined *LacZ/orJ* mutant with the *mi* mutation, the retinal phenotype was not characterized as part of this study. *Mi* mutant mice with the *LacZ* allele in a homozygous state would be needed to compare the effects of reduced Mitf activity on the *LacZ* mutant phenotype compared to the *orJ* mutant phenotype.

In summary, the *LacZ* allele is a new addition to the allelic series of *Vsx2* mutations, and its functional properties make it useful for further probing *Vsx2* function in the *in vivo* context.

## Experimental Procedures

### Animals

The *Vsx2^LacZ^* line was generated from ES cells containing the *Vsx2^tm1a(EUCOMM)Wtsi^* allele. Germline founders were crossed with CMV-Cre mice (RRID:IMSR_JAX:006054) to excise the *LoxP* flanked region containing the Neo cassette and *Vsx2* exon 3 to create the *Vsx2^LacZ^* strain. *Vsx2^orJ^* mice (RRID:IMSR_JAX:000395) and *Mitf^mi^* mice (RRID:IMSR_JAX001573) are maintained in the laboratory.

### Tissue collection

Embryos and pups were generated using single night matings, with the day that the vaginal plug was observed considered E0.5. *Vsx2^LacZ^* embryos were collected at E10.5, E11.5, E12.5, E13.5, E15.5 and *Vsx2^LacZ^* pups were collected at P0, P10, P14, and P28. All embryos and pups were staged using whole-body attributes.^23–25^

To assess proliferation, EdU was delivered via intra-peritoneal injection at a dose of 30 µg/g body weight at two hours prior to euthanasia of dam for embryonic dissection and 45 minutes prior to euthanasia for postnatal pups. Dams and pups from P10 and older were euthanized using CO_2_ and cervical dislocation; P0 pups were anesthetized using hypothermia and thereafter euthanized via decapitation. At E15.5, embryo eyelids were removed for imaging to allow for accurate eye size quantification.

### Tissue preparation

Embryo heads and postnatal pup eyes were fixed at room temperature in 4% PFA-PBS solution for 45 minutes or 1 hour, respectively. After several PBS washes, a sucrose cryopreservation series was performed before the samples were frozen in TissueTek OCT (Sakura Finetech) and stored at −80°C. Tissue was sectioned in twelve micrometer sections at −20°C with a Leica CM1950 cryostat and adhered to Superfrost Plus slides (Fisher Scientific). The sections were then dried for 1-2 hours before storage at −20°C and −80°C.

### PCR

DNA was extracted from embryonic tissue samples by adding 75µL alkaline lysis buffer to sample tube, heating to 95°C for 10 minutes, cooling for 10 minutes, followed by adding 75µL of neutralization buffer. DNA extractions were stored at −20°C.

In a 10:1 ratio, 9µL of master mix was added to 1µL extracted DNA and run in a ProFlex thermocycler (Life Technologies).

A 1.5% agarose gel was used to run the PCR product alongside a Quick-Load 1kb Plus DNA Ladder at 70 V for 95 minutes. A UVP Gel Doc-It^TS3^ Imager was used to image the gel.

The PCR and gel electrophoresis confirming germline transmission of the LacZ allele is a representative image from three replicates.

### Western Blot

Freshly dissected E15.5 retinal tissue was flash frozen in liquid nitrogen and stored at −80°C. Lysates were prepared with RIPA buffer and EDTA-free protease inhibitor (Millipore Sigma) at a volume of 100 µL per 50 mg of tissue. Lysates for heterozygotes contained 2 retinas per sample (1 animal) and 4 retinas per sample (2 animals) for homozygotes. BCA assay (Quick Start Bradford, Bio rad) was used to determine protein concentration using a wavelength of 595 nm. 20 ug of protein per lane was loaded in a 20 µL volume. Gels were run for 40 min at 90 V and then 40 min at 110 V. Proteins were transferred using an iBlot2 (Fisher Scientific) system with a setting of 15V in 7 mins. Blots were incubated for 1 hour at room temperature in 1X PBS, pH 7.4, 0.1% Tween 20 containing 5% bovine serum albumin (PBST/B), followed by incubation with constant rocking overnight at 4°C with primary antibody diluted in PBST/B (Table 1). Excess antibody was removed, and blots were washed 3 times for 10 minutes each with PBST/B, then incubated for 1 hour at room temperature with secondary antibody diluted in PBST (Table 1). Excess antibody was removed and blots were washed 6 times for 10 minutes each with PBST. Chemiluminescence HRP substrate (Immobilon Western, Millipore) was used for signal detection and images were captured on an Amersham Imager 600 (General Electric). The western blot presented is a representative image from 3 replicates.

### Immunohistochemistry and EdU detection

To remove OCT, slides were washed in PBS and incubated in blocking solution (PBST (0.05% Tween 20) containing 2% normal donkey serum (PBST/DS)) for 1 hour at room temperature. Slides were then incubated overnight at 4°C in primary antibody diluted in PBST/DS (Table 1). Excess antibody was removed, slides were washed 3 times for 5 minutes each with PBST, followed by incubation for 1 hour at room temperature in a dark humidified chamber with secondary antibodies diluted in PBST/DS (Table 1). Excess antibody was removed, and slides were washed 3 times for 10 minutes each with PBST. EdU incorporation was detected using AlexaFluor 568 or 647 Click-iT Cell Reaction Kit (Invitrogen-Molecular Probes, Eugene, Oregon). DAPI (1:40000 in PBS) was used to label nuclei. After final washes in dH_2_O, slides were air dried and coverslipped with Fluoromount and stored at 4°C until imaging. All immunohistology expression patterns are representative images from a sample size of more than three, collected from at least two different litters for each stage and genotype.

### Quantifications

Eye size was quantified using the pentagon-ROI tool of Fiji on embryo pictures taken on a Nikon C-FIDH SMZ1270. Eye size was defined by the area of the two-dimensional circumference of the pigmented eye (sphere). Area was measured in pixels^2^ and converted to µm^2^. Neurogenic length, as defined by the linear length of the retinal basal surface that TUBB3-positive cells occupy, was quantified using the freehand line-ROI tool of Fiji and the percentage calculated by dividing the neurogenic length by the total length of the retinal basal surface and multiplying by 100. Neurogenic density, as defined by the number of TUBB3-positive pixels divided by the total number of pixels in the neurogenic region (the apicobasal region of the retina containing TUBB3-positive cells), is used as a proxy for cells within the neurogenic region. This was quantified using Fiji by creating an ROI outlining the TUBB3-positive region of the retina and clearing the non-ROI regions using Edit>Clear Outside. The neurogenic region was then made binary using Threshold tool, then the number of positive (white) and negative (black) pixels were quantified using Analyze>Histogram. The neurogenic density was then calculated using the ratio of positive pixels to total pixels in the neurogenic region.

### Statistical Analysis

Prism was used to run all statistical analyses and to create graphical representations. Unpaired two-tailed T-tests were used for comparisons limited to 2 conditions (Fig. 1L; Fig. 5B,H,I; Fig. 6C,F). The coefficient of variation within a sample group was calculated using the formula CV=(Standard Deviation / Sample Mean)*100 (Fig. 5C,J,K). One way ANOVAs accompanied by post-hoc analyses (Brown-Forsythe, Bartlett’s, Tukey’s multiple comparison tests) were run using Prism on the E15.5 eye size, neurogenic length, and neurogenic density data to thoroughly compare the heterozygous and homozygous phenotypes of the *LacZ* and *orJ* alleles (Fig. 5B,H,I). Linear regression was used to describe the relationship between eye size and measures of neurogenesis (neurogenic density (Fig. 5L) and neurogenic length (Fig. 5M). Supplemental Table 1 lists the results of all statistical analyses.

### Imaging/Microscopy

Whole embryos were imaged using Nikon C-FIDH SMZ1270. Immunofluorescent staining at E10.5-E11.5 and postnatal stages were imaged using a Nikon Eclipse E600 while those at E12.5-E15.5 were imaged using a Zeiss LSM710 confocal microscope with 20X lens to capture a high-resolution image of the relatively larger ROI. Image histograms were adjusted using Fiji. Images were resized and trimmed using Adobe Photoshop. Figures were made using Adobe Illustrator.

## Acknowledgments

Funding was provided by the National Eye Institute (R01-EY013760; P30-EY008126) and an unrestricted grant from Research to Prevent Blindness, Inc. AAH was supported by the National Science Foundation Graduate Research Fellows Program (1937963). The Zeiss LSM710 confocal microscope used for this study was acquired with funding from NIH (S10-RR027396). We thank members of the Levine and Fuhrmann laboratories for their insights and feedback.

## Ethics Statement

All procedures and experiments with mice were approved by the Vanderbilt Institutional Animal Care and Use Committee and conformed to the ARVO guidelines for the use of animals in vision research. The research was conducted in concordance with the NIH guidelines for Responsible Conduct in Research.

## Disclosures

The authors report no conflicts of interest in this work.

---

**Supplemental Figure 1. Expression of VSX2 and b-GAL in postnatal *LacZ* het retina. (A,B)** Immunohistology reveals b-GAL to be an accurate reporter of VSX2 expression with both expressed throughout the *LacZ* het retina at P0. **(C-F)** At P10 and P14, VSX2 and b-GAL are restricted to the INL, consistent with expression in bipolar cells and Müller glia. All VSX2 and b-GAL stains are from the same animal for each respective stage and genotype. Scale bar: 100 µm.

**Supplemental Figure 2. Immunohistology at E10.5 and E13.5. (A-D)** Expression of the proliferation markers PCNA, CCND1, and pHH3 in the *LacZ* mutant retina are normal at E10.5. **(E-J)** At E13.5, PCNA extends throughout the *LacZ* mutant retina (E,F), whereas CCND1 and EdU are limited to the central retina (G-J). Arrows in G-J point to areas where staining is absent in the *LacZ* mutant compared to the *LacZ* het. **(K,L)** ATOH7 expression is detected in a few cells in the central retina of the *LacZ* mutant (arrow). **(M-P)** Retinal ganglion cells (POU4F2) and amacrine cells (BHLHB5) are not detected in the *LacZ* mutant retina at E13.5. **(Q,R)** OTX1/2 expression is detected in a few cells in the central retina of the *LacZ* mutant (arrow). **(S,T)** SOX2 expression is reduced in the peripheral retina of the *LacZ* mutant. Arrows in S and T point to areas where staining is absent in the *LacZ* mutant compared to the *LacZ* het. Scale bars: 100 µm.

**Supplemental Figure 3. *LacZ* mutant retina at P14. (A-F)** As the P14 retina continues to aberrantly proliferate (EdU, MCM6) and give rise to new neurons (TUBB3), signs of degeneration emerge. **(C-P)** The previously organized apical epithelium composed largely of progenitors (MCM6, SOX9), Müller glia (SOX9), and photoreceptors (OTX1/2, RECOVERIN) begins to seemingly lose its integrity as amongst the disorganized basal cell population, consisting of progenitors (MCM6, SOX9), Müller glia (SOX9), retinal ganglion cells (POU4F2) and amacrine cells (BHLHB5), hypocellular regions, possibly plexiform outgrowths, encroach. **(Q,R)** PKC expressing cells are present in the *LacZ* mutant retina, but their identity is not known, with possibilities being amacrine cells or remnant bipolar cells. **(S,T)** VSX1 expressing cells are not detected in the *LacZ* mutant retina. Scale Bar: 100 µm.

**Supplemental Figure 4. Uncropped images for western blots in Figure 1C. (A)** Blot 1 probed with N-terminal VSX2 antibody, which detects the full length (VSX2 WT) and truncated (VSX2 Mut) proteins. The last 4 lanes in panel A were used for the first two rows in Fig. 1C. **(B)** Blot 1 stripped and re-probed with antibody to a validated reference protein ATP5A1. The last 4 lanes in panel B were used for the third row in Fig. 1C. **(C)** Blot 2 probed with b-GAL antibody. Spacings between lanes were cropped and the bands at 100 kDa correspond to the fourth row in Fig. 1C. **(D)** Blot 2 was stripped and reprobed with ATP5A1 antibody. Spacings between lanes were cropped and the bands at 50 kDa correspond to the fifth row in Fig. 1C.

## Notes

### Competing Interest Statement

The authors have declared no competing interest.

### Summary of Updates

Sample sizes, p-values, and statistical tests are provided in Figure legends. Supplemental Table 1 provides statistical summaries for all quantifications. Table 2 lists secondary antibodies and their corresponding dilutions. Methods have been updated to provide more clarity for reproducibility. The discussion now includes a section describing the limitations of the study.

## References

1. Burmeister M, Novak J, Liang MY, et al. Ocular retardation mouse caused by Chx10 homeobox null allele: impaired retinal progenitor proliferation and bipolar cell differentiation. Nat Genet. 1996;12(4):376–384. doi:10.1038/ng0496-376

2. Liu ISC, Chen J de, Ploder L, et al. Developmental expression of a novel murine homeobox gene (Chx10): Evidence for roles in determination of the neuroretina and inner nuclear layer. Neuron. 1994;13(2):377–393. doi:10.1016/0896-6273(94)90354-9

3. Zou C, Levine EM. Vsx2 Controls Eye Organogenesis and Retinal Progenitor Identity Via Homeodomain and Non-Homeodomain Residues Required for High Affinity DNA Binding. Desplan C, ed. Plos Genet. 2012;8(9):e1002924. doi:10.1371/journal.pgen.1002924

4. Phillips MJ, Perez ET, Martin JM, et al. Modeling human retinal development with patient-specific iPS cells reveals multiple roles for VSX2. Stem Cells. Published online 2014:n/a-n/a. doi:10.1002/stem.1667

5. Theiler K, Varnum DS, Nadeau JH, Stevens LC, Cagianut B. A new allele of ocular retardation: early development and morphogenetic cell death. Anat Embryol. 1976;150(1):85–97. doi:10.1007/bf00346288

6. Green ES, Stubbs JL, Levine EM. Genetic rescue of cell number in a mouse model of microphthalmia: interactions between Chx10 and G1-phase cell cycle regulators. Development. 2003;130(3):539–552. doi:10.1242/dev.00275

7. Sigulinsky CL, German ML, Leung AM, Clark AM, Yun S, Levine EM. Genetic chimeras reveal the autonomy requirements for Vsx2 in embryonic retinal progenitor cells. Neural Dev. 2015;10(1):12. doi:10.1186/s13064-015-0039-5

8. Leung AM, Rao MB, Raju N, et al. A framework to identify functional interactors that contribute to disrupted early retinal development in Vsx2 ocular retardation J mice. Dev Dyn. Published online 2023. doi:10.1002/dvdy.629

9. Horsford DJ, Nguyen MTT, Sellar GC, Kothary R, Arnheiter H, McInnes RR. Chx10 repression of Mitf is required for the maintenance of mammalian neuroretinal identity. Development. 2005;132(1):177–187. doi:10.1242/dev.01571

10. Rowan S, Chen CMAM, Young TL, Fisher DE, Cepko CL. Transdifferentiation of the retina into pigmented cells in ocular retardation mice defines a new function of the homeodomain gene Chx10. Development. 2004;131(20). doi:10.1242/dev.01300

11. Bone Larson C, Basu S, Radel JD, et al. Partial rescue of the ocular retardation phenotype by genetic modifiers. J Neurobiol. 2000;42(2):232–247. doi:10.1002/(sici)1097-4695(20000205)42:2<232::aid-neu7>3.0.co;2-4

12. Bharti K, Liu W, Csermely T, Bertuzzi S, Arnheiter H. Alternative promoter use in eye development: the complex role and regulation of the transcription factor MITF. Development. 2008;135(6):1169–1178. doi:10.1242/dev.014142

13. Percin EF, Ploder LA, Yu JJ, et al. Human microphthalmia associated with mutations in the retinal homeobox gene CHX10. Nat Genet. 2000;25(4):397–401. doi:10.1038/78071

14. Kurtzman AL, Schechter N. Ubc9 interacts with a nuclear localization signal and mediates nuclear localization of the paired-like homeobox protein Vsx-1 independent of SUMO-1 modification. Proc Natl Acad Sci USA. 2001;98(10):5602–5607. doi:10.1073/pnas.101129698

15. Dorval KM, Bobechko BP, Ahmad KF, Bremner R. Transcriptional activity of the paired-like homeodomain proteins CHX10 and VSX1. J Biol Chem. 2005;280(11):10100–10108. doi:10.1074/jbc.m412676200

16. Skarnes WC, Rosen B, West AP, et al. A conditional knockout resource for the genome-wide study of mouse gene function. Nature. 2011;474(7351):337–342. doi:10.1038/nature10163

17. Knauer SK, Carra G, Stauber RH. Nuclear Export Is Evolutionarily Conserved in CVC Paired-Like Homeobox Proteins and Influences Protein Stability, Transcriptional Activation, and Extracellular Secretion. Mol Cell Biol. 2005;25(7):2573–2582. doi:10.1128/mcb.25.7.2573-2582.2005

18. Konyukhov B, Sazhina M. Interaction of the genes of ocular retardation and microphthalmia in mice. Folia Biol (Praha). 1966;12(2):116–123. http://eutils.ncbi.nlm.nih.gov/entrez/eutils/elink.fcgi?dbfrom=pubmed&id=4958088&retmode=ref&cmd=prlinks

19. Goodson NB, Kaufman MA, Park KU, Brzezinski JA. Simultaneous deletion of Prdm1 and Vsx2 enhancers in the retina alters photoreceptor and bipolar cell fate specification, yet differs from deleting both genes. Development. 2020;147(13):dev190272. doi:10.1242/dev.190272

20. Wong G, Conger SB, Burmeister M. Mapping of genetic modifiers affecting the eye phenotype of ocular retardation (Chx10or-J) mice. Mamm Genome. 2006;17(6):518–525. doi:10.1007/s00335-005-0159-z

21. Rowan S, Cepko CL. Genetic analysis of the homeodomain transcription factor Chx10 in the retina using a novel multifunctional BAC transgenic mouse reporter. Dev Biol. 2004;271(2):388–402. doi:10.1016/j.ydbio.2004.03.039

22. Kurtzman AL, Gregori L, Haas AL, Schechter N. Ubiquitination and degradation of the zebrafish paired-like homeobox protein VSX-1. J Neurochem. 2000;75(1):48–55. doi:10.1046/j.1471-4159.2000.0750048.x

23. Wong MD, Eede MC van, Spring S, et al. 4D atlas of the mouse embryo for precise morphological staging. Development. 2015;142(20):3583–3591. doi:10.1242/dev.125872

24. Theiler K. The House Mouse: Atlas of Embryonic Development. Springer, Berlin; 1989. doi:10.1007/978-3-642-88418-4

25. Kaufman MH. The Atlas of Mouse Development. Elsevier Science; 1992. https://books.google.com/books?id=Lo5pAAAAMAAJ

26. Clark AM, Yun S, Veien ES, et al. Negative regulation of Vsx1 by its paralog Chx10/Vsx2 is conserved in the vertebrate retina. Brain Res. 2008;1192:99–113. doi:10.1016/j.brainres.2007.06.007

