## Supplementary figures and images for "Microphthalmia and disrupted retinal development due to a *LacZ* knock-in/knock-out allele at the *Vsx2* locus"

### Supplemental Figure 1

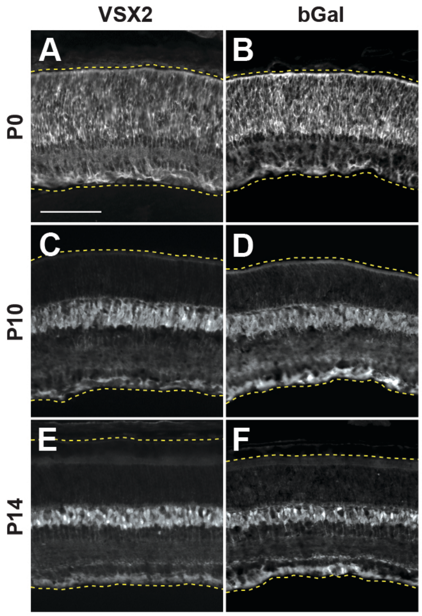


All scale bars = 100um

### Supplemental Figure 2

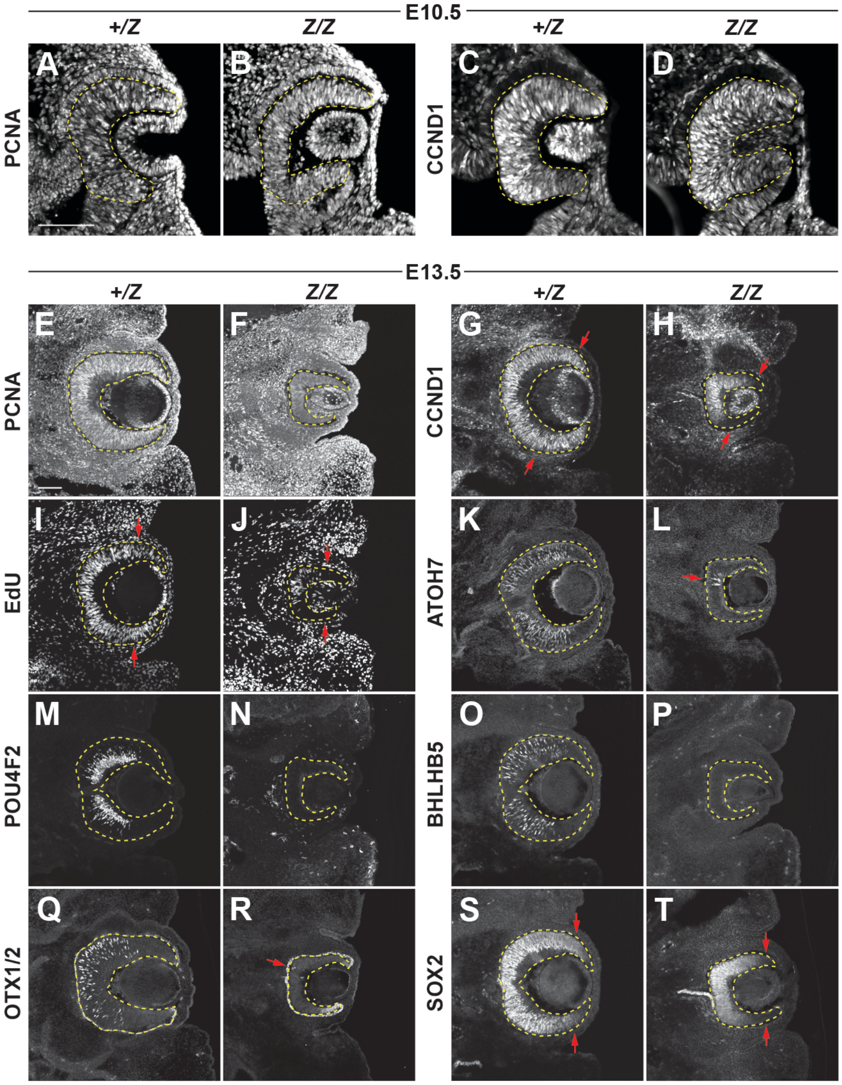

### Supplemental Figure 4

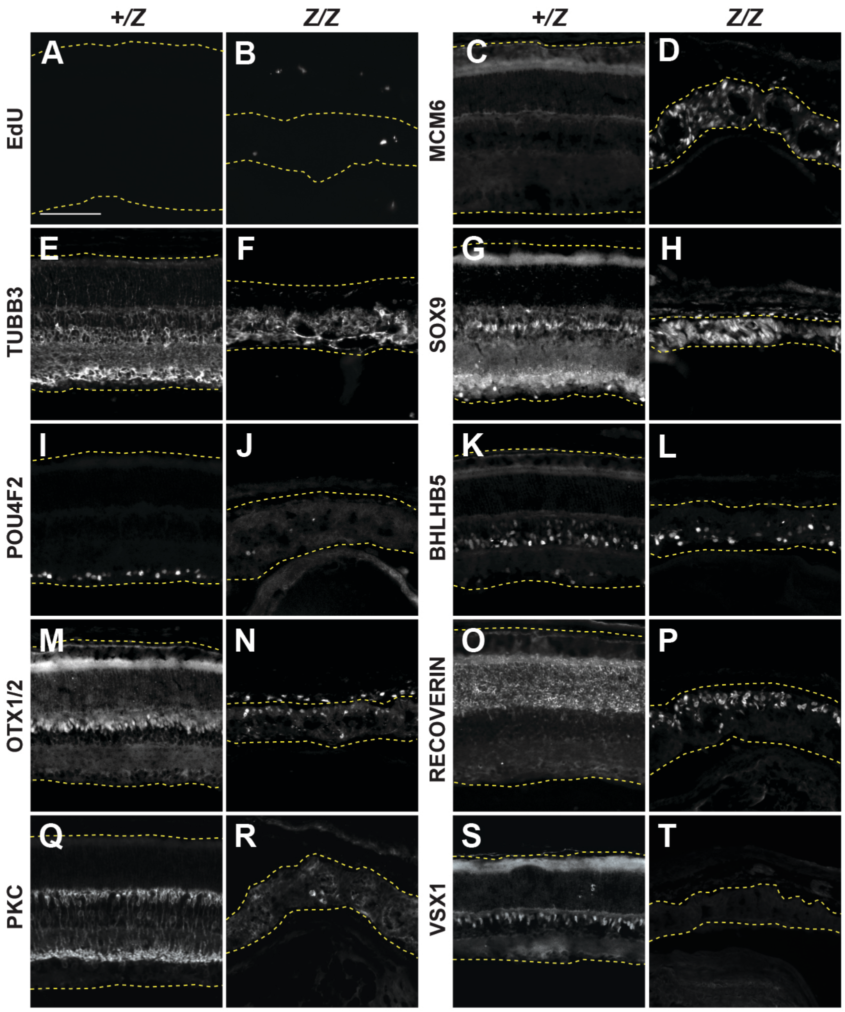

### Supplemental Figure 4

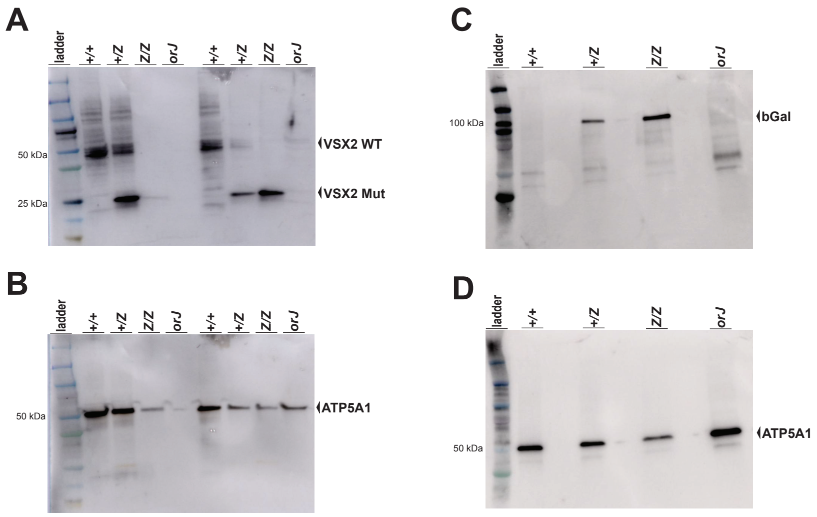
